# Reprogramming LCLs to iPSCs Results in Recovery of Donor-Specific Gene Expression Signature

**DOI:** 10.1101/013631

**Authors:** Samantha M. Thomas, Courtney Kagan, Bryan J. Pavlovic, Jonathan Burnett, Kristen Patterson, Jonathan K. Pritchard, Yoav Gilad

## Abstract

Renewable *in vitro* cell cultures, such as lymphoblastoid cell lines (LCLs), have facilitated studies that contributed to our understanding of genetic influence on human traits. However, the degree to which cell lines faithfully maintain differences in donor-specific phenotypes is still debated. We have previously reported that standard cell line maintenance practice results in a loss of donor-specific gene expression signatures in LCLs. An alternative to the LCL model is the induced pluripotent stem cell (iPSC) system, which carries the potential to model tissue-specific physiology through the use of differentiation protocols. Still, existing LCL banks represent an important source of starting material for iPSC generation, and it is possible that the disruptions in gene regulation associated with long-term LCL maintenance could persist through the reprogramming process. To address this concern, we studied the effect of reprogramming mature LCLs to iPSCs on the ensuing gene expression patterns within and between six unrelated donor individuals. We show that the reprogramming process results in a recovery of donor-specific gene regulatory signatures. Since environmental contributions are unlikely to be a source of individual variation in our system of highly passaged cultured cell lines, our observations suggest that the effect of genotype on gene regulation is more pronounced in the iPSCs than in the LCL precursors. Our findings indicate that iPSCs can be a powerful model system for studies of phenotypic variation across individuals in general, and the genetic association with variation in gene regulation in particular. We further conclude that LCLs are an appropriate starting material for iPSC generation.

## Introduction

Renewable cell models are widely recognized as valuable platforms for studies of human genotype-phenotype interactions because they are easily manipulated, scalable, and are specific to human physiology (in contrast to lab animal models). Epstein-Barr virus (EBV) transformed lymphoblastoid cell lines (LCLs) are one such commonly-used model. In recent years, LCLs have been used to study genetic influence on disease traits [1], drug response [2-5], and gene regulation [6,7]. In particular, much of what we now know about associations of human genetic variation with differences in gene regulation is based on studies that used data from LCLs. There is little doubt that many fundamental regulatory principles that we have learned by generating and analyzing data from LCLs are generally shared with primary tissues. However, a critical property of any *in vitro* cellular model is the ability to faithfully recapitulate the specific regulatory properties of the donor’s primary tissue. In that regard, though LCLs have clearly been a convenient and useful model, there is concern that factors related to immortalization and cell line maintenance obscure genetic signal in LCLs [8-10].

A number of studies have characterized differences in gene regulatory phenotypes between LCLs and primary tissues [11-14]. These have shown that a large number of genes are differentially expressed between primary cells and cell lines, and that thousands of CpG sites are differentially methylated between LCLs and primary blood cells. Our group has also demonstrated disruptions in gene regulation in LCLs by studying multiple independent replicates of LCLs from isolated primary B cells of six individuals and repeatedly subjecting the cell lines to cycles of freeze, thaw, and recovery. We found that newly transformed LCLs (within a few passages after the EBV transformation) largely maintained individual differences in gene expression levels. However, LCLs that had been frozen and thawed at least once (we referred to these as mature LCLs) exhibited a substantial loss of inter-individual variation in gene expression levels [14,15].

On the one hand, it is unlikely that the loss of the donor effect on gene expression would lead to false positive findings of genetic influence on gene regulation. Indeed, we reported that genes associated with previously identified eQTLs retain relatively high variation in gene expression levels between individuals even after repeated freeze-thaw culturing cycles. Yet on the other hand, because much of the individual variation observed in primary tissues is not exhibited by LCLs, studies using the LCL model are limited in their ability to detect donor differences.

The induced pluripotent stem cell (iPSC) system is another renewable cell model that is increasingly used to study individual phenotypic variation because it can ultimately provide access to a wide range of tissue types through the use of differentiation protocols. However, the capacity of iPSCs and derived cell types to faithfully recapitulate *in vivo* physiology is also still largely unknown. Previous studies have noted a significant effect of donor on traits in iPSCs such as hematopoietic [16], neuronal [17] and hepatic [18] differentiation potential. Importantly, the genetic background of iPSCs generated from peripheral blood mononuclear cells and fibroblasts was recently demonstrated to account for more of the variation in gene expression between iPSC lines than any other tested factor such as cell type of origin or reprogramming method [19]. While these findings indicate that reprogramming iPSCs from primary tissues preserves individual variation in gene expression, it is unknown whether reprogramming highly manipulated immortalized cell lines, such as LCLs, to iPSCs can recover the individual gene expression patterns lost during cell line maintenance.

Because LCLs are available in large banks representing disease populations or ethnicities, they are a promising source of starting material for iPSC generation if disruptions in gene regulation do not persist through the reprogramming process. In the present study, we ask whether reprogramming mature LCLs to iPSCs can result in the recovery of individual variation in gene expression that had been lost during the LCL maturation and maintenance process.

## Results

To test whether reprogramming LCLs to iPSCs could recover the effect of donor on gene expression profiles, we generated iPSCs from three mature LCLs of each of six Caucasian individuals for a total of 17 pairs of cell lines (one iPSC line failed to reach the requisite ten passages and was excluded from the study; see methods). We have previously collected gene expression data from the LCLs at earlier stages [14,15]. For the current study, we quantified whole genome gene expression microarray data from the 17 mature LCLs immediately prior to reprogramming and from stable and validated iPSCs. See Fig. 1 for schematic of the study design and S1 Table for the processed gene expression data from all samples.

**Figure 1.**
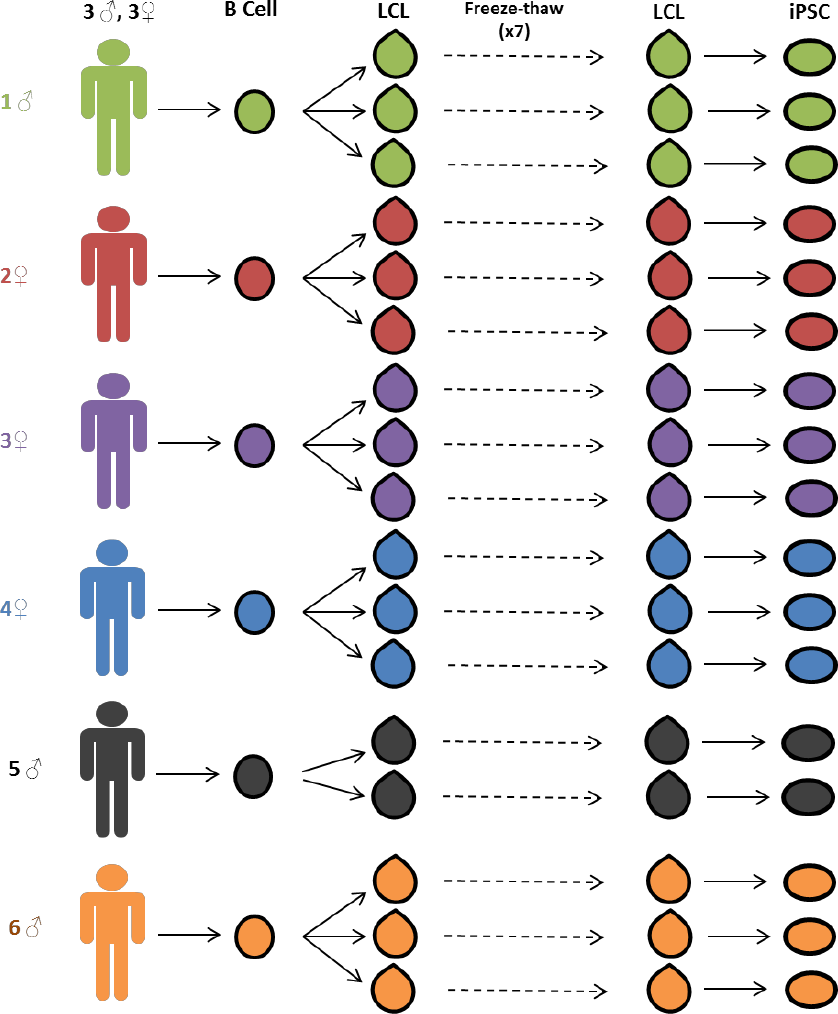
Study design. Three independent lymphoblastoid cell lines (LCLs) were generated for each of six unrelated Caucasian individuals. LCLs were frozen and thawed seven times. After the seventh thaw, the LCLs were reprogrammed to iPSCs. Gene expression data was collected from LCLs immediately before reprogramming and from stable iPSC lines.

### Generation and Validation of the iPSCs

We reprogrammed mature LCLs, which had previously undergone seven freeze-thaw culturing cycles, to iPSCs using an episomal transfection approach [20-22] (see Methods for more details). We reprogrammed the LCLs in four batches; scheduling LCLs derived from the same individual to different reprograming batches to ensure that no artificial correlation structure was introduced between ‘reprograming batch’ and ‘donor individual’ in the process of iPSC generation. All iPSC lines were confirmed to be pluripotent using an embryoid body assay (Fig. 2A and S1 Fig.), qPCR for pluripotency-associated transcription factors (Fig. 2B), genomic PCR to confirm the absence of reprogramming plasmids (S2 and S3 Figs.), and PluriTest [23] (S2 Table). Three independently established LCLs were successfully reprogrammed into validated iPSCs for all but one individual, for which only two iPSC lines were obtained.

**Figure 2.**
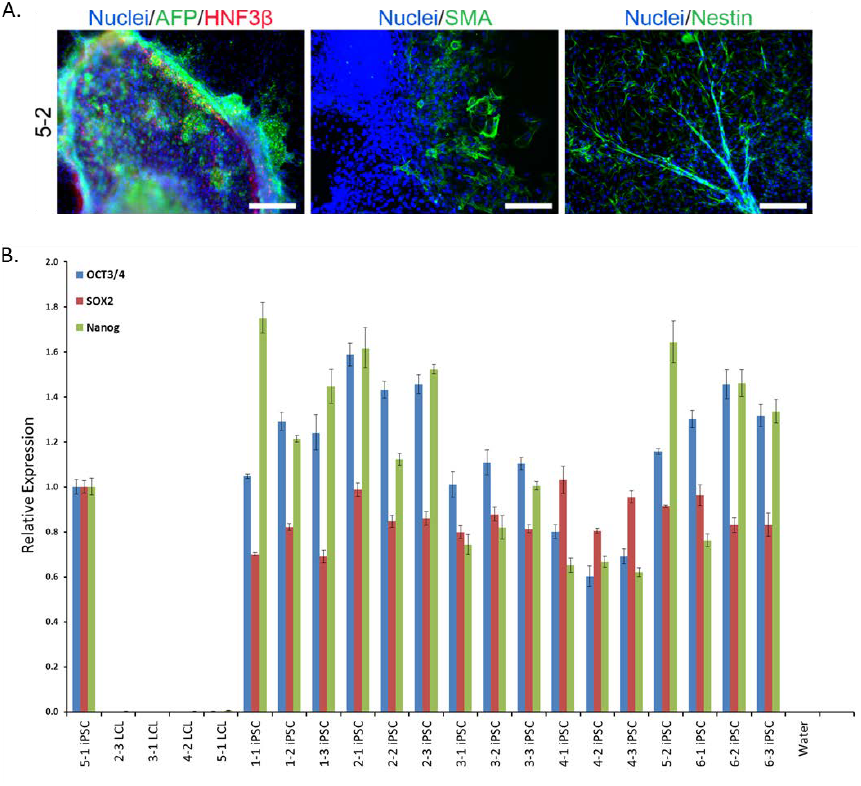
iPSC generation and validation. A. Representative embryoid body staining for iPSC line 5-2 demonstrating differentiation potential for endoderm, mesoderm, and ectoderm lineages. See S1 Fig. for results from all lines. Scale bars represent 200 μm B. Results from qPCR for three endogenous pluripotency-related transcription factors, normalized to GAPDH. iPSC 5-1 was randomly chosen as a reference sample.

### Recovery of the Individual Signature of Gene Regulation

We collected high quality RNA (RIN score range: 7.6-9.9; S2 Table) from LCLs immediately prior to reprogramming and from the stable and validated iPSC lines after at least 10 passages (see S2 Table for specific passage information). We quantified gene expression levels for all samples using the Illumina Human HT12v4 microarray platform. As a first step of our analysis, we excluded data from probes whose target transcripts did not map to a unique Ensembl gene ID and those that were not detected as ‘expressed’ in at least two samples from either cell type (we note that our general observations are robust with respect to a wide range of this inclusion criteria). We also excluded from the analysis data from probes with a known HapMap SNP with a minor allele frequency > 0.01 in the CEU population, to eliminate the possibility of an artificial effect of genotype on the hybridization-based estimates of gene expression levels. We then quantile-normalized the combined data from the remaining probes across all samples. We examined and corrected for array batch using the approach of Johnson et al [24] (see Methods). Finally, we obtained normalized expression levels for 15,306 genes detected as expressed in our samples (S1 Table). Using a linear model-based Empirical Bayes method (implemented in the ‘*limma*’ R package [25]), we classified 9,746 genes as differentially expressed between iPSCs and LCLs (FDR < 1%; see Methods for more details about modeling and hypothesis testing).

Because the regulation of a large percentage of genes was affected by reprogramming (64% of tested genes), we asked whether gene expression patterns specific to the donor individual were recovered in the process. We addressed this question using two approaches. First, we evaluated the overall degree of similarity across cell lines from the same donor by considering summaries of the gene expression phenotypes using clustering analysis and PCA. The rationale for collapsing our gene-specific expression data and considering overall summaries is that complex phenotypes can often be the result of a large combination of genotype contributions and we are interested to learn whether the overall data from cell lines exhibits a clear signature of the donor. In our second approach, we focused on gene specific patterns by partitioning the variance in expression levels for individual genes and testing for differences between the entire distributions of gene expression levels across cell lines. In this approach we are considering expression patterns of individual genes as independent data points. The rationale for the gene-specific approach is that studies of the genetic basis for regulatory variation (such as eQTL mapping studies) nearly always consider the expression phenotypes of individual genes and we are interested to learn the extent to which the effect of donor genotype on gene expression levels can be studied using a given cell model.

To evaluate overall clustering properties in the expression data from the two cell types, we performed hierarchical clustering analysis and PCA. As we performed these analyses, we consistently observed that data from the second iPSC line of individual 4 (line marked as 4-2 in our figures) accounts for a disproportionate amount of variance (S4B Fig.). This individual is a clear outlier and its iPSC is associated with the lowest PluriScore in our study (S2 Table). We have excluded the data from this individual from subsequent analyses. Importantly, we have confirmed (as we show in supplementary figures), that our conclusions our robust with respect to this decision.

Using data from all 15,306 genes detected as expressed, mature LCLs fail to consistently cluster by the individual from whom they were initially derived, in accordance with our previous observations (Fig. 3A, S4A/5A Figs.). Data from the corresponding iPSC lines, however, cluster by the individual of origin, indicating a large degree of recovery of donor gene expression patterns (Fig. 3B, S4B/5B Figs.). Another method to assess overall clustering properties is through the use of principal components analysis. Taking this approach, we found that clustering of the expression data by individual of origin is substantially more pronounced in the iPSCs than in the LCLs (Fig 3 and S4 Fig.). Indeed, the average pairwise Euclidean distances of expression data projections on the first two PCs are significantly smaller within cell lines derived from the same individual than those from different individuals for iPSCs (*P* < 10^-12^), but not for LCLs (*P* = 0.10; S3 Table).

**Figure 3.**
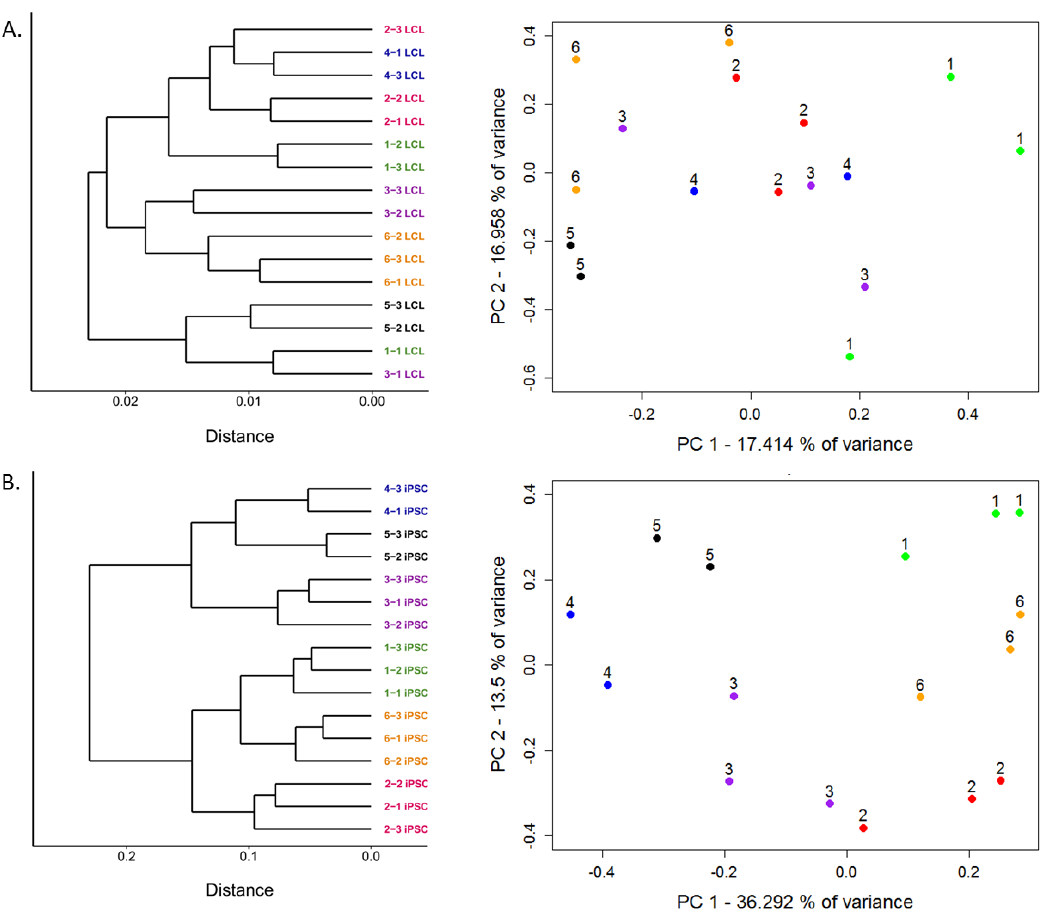
Improved clustering properties after reprogramming to iPSCs. A. Results from hierarchical clustering analysis of microarray gene expression and expression data projections on principal components axes 1 and 2 from cycle 7 LCLs and B. iPSCs.

Turning our attention to expression patterns of individual genes, we estimated the magnitude of the donor effect on gene expression patterns in LCLs and iPSCs. To do so, we compared the pairwise correlations of expression data from cell lines derived from the same donor to pairwise correlations of data from cell lines derived from different individuals (S6 Fig.). On average, both within- and between-donor correlation coefficients are significantly higher in iPSCs than in the LCLs they were initially derived from (p < 10^-4^ and p < 10^-7^ for within- and between-donor correlations, respectively). In other words, regardless of the individual of origin, we observed less variation in gene expression between iPSCs than LCLs. Yet, though iPSCs harbor less variation overall, the proportion of variation in gene expression that is explained by donor is significantly higher in the iPSCs compared with the LCLs (*P* < 10^-15^). Indeed, using a single factor ANOVA, we estimate that donor explains, on average, 23.4% of the variance in gene expression in iPSCs but only 6.6% in LCLs.

In addition to within-donor correlations, we were specifically interested in identifying genes that were highly variable across donors. We thus proceeded by considering the ratio of between-to within-individual variation in gene expression levels in the two cell types. On average, we found a significantly higher ratio of between-to-within individual variance in gene expression levels in iPSCs compared with data from the LCLs (*P* < 10^-15^; recall that in this analysis we consider expression patterns of individual genes as independent data points), despite significantly higher overall variance in LCL gene expression (*P* < 10^-9^; Fig. 4, S7 Fig.). We identified 1,831 genes whose expression levels were significantly associated with donor in iPSCs (single factor ANOVA FDR < 0.05; see S8 Fig. for histogram of p-values) but only 104 such genes in LCLs.

**Figure 4.**
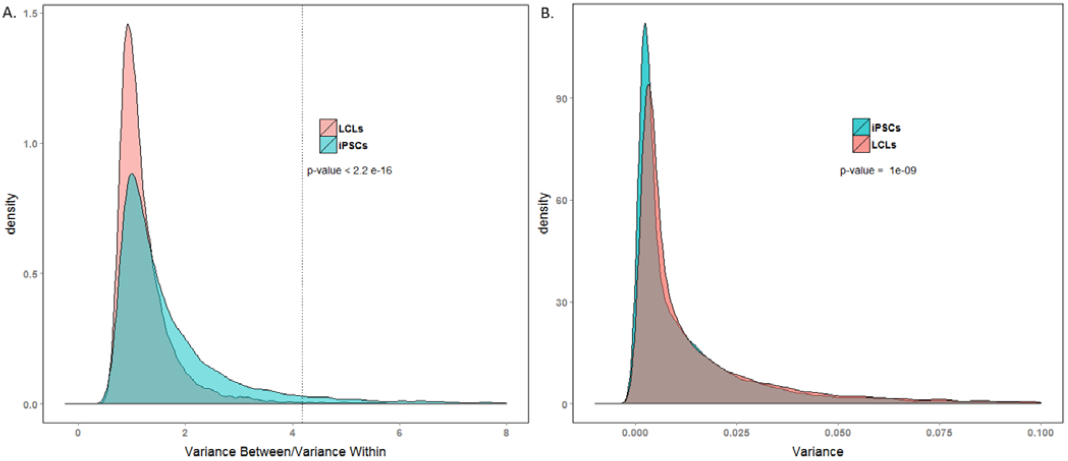
Comparison of ability to detect inter-individual gene expression variation. Density plot of between donor variance to within donor variance in gene expression for all expressed genes in iPSCs and LCLs. The dotted line indicates the threshold ratio corresponding with significant association between gene expression and donor. X-axis was truncated at 8.0; 0.77% of the data are not shown. B. Density plot of total variance in LCLs and iPSCs. X-axis was truncated at 0.1; 3.4% of the data are not shown.

### Functional Relevance of Highly Variable Genes

We tested for enrichment of functional annotation related to tissue-expression (using the online database Lynx [26]) among genes whose expression levels are significantly associated with donor. While these results do not shed much light on the functional importance of these gene sets, we note that different classes of genes exhibit high individual variation in the two cell types. For example, genes with a strong donor effect in LCLs are enriched with genes expressed in blood and those in iPSCs are enriched in genes expressed in embryonic tissue. The complete set of enrichment results is available in S4 and S5 Tables.

Finally, we considered the relevance of our findings with respect to previously published eQTL studies in LCLs. Our sample of 6 individuals is too small to allow identification of eQTLs. As an alternative, we compared individual variation in expression levels between genes previously associated with an eQTL in LCLs [6], and genes for which an eQTL was not identified. To do so, we randomly selected data from one biological replicate (one LCL and its corresponding iPSC) from each individual.

In both LCLs and iPSCs, the average coefficients of expression variation were significantly higher in genes previously associated with eQTLs than in genes for which eQTLs were not identified (*P* < 10^-10^ and *P* = 0.002, for LCLs and iPSCs, respectively; Fig. 5, S9 Fig.). As expected (given that these eQTLs were originally observed in LCLs, and that LCLs have greater overall variation), the coefficients of variation are significantly higher in eQTL-associated genes in LCLs than iPSCs (*P* < 10^-8^).

**Figure 5.**
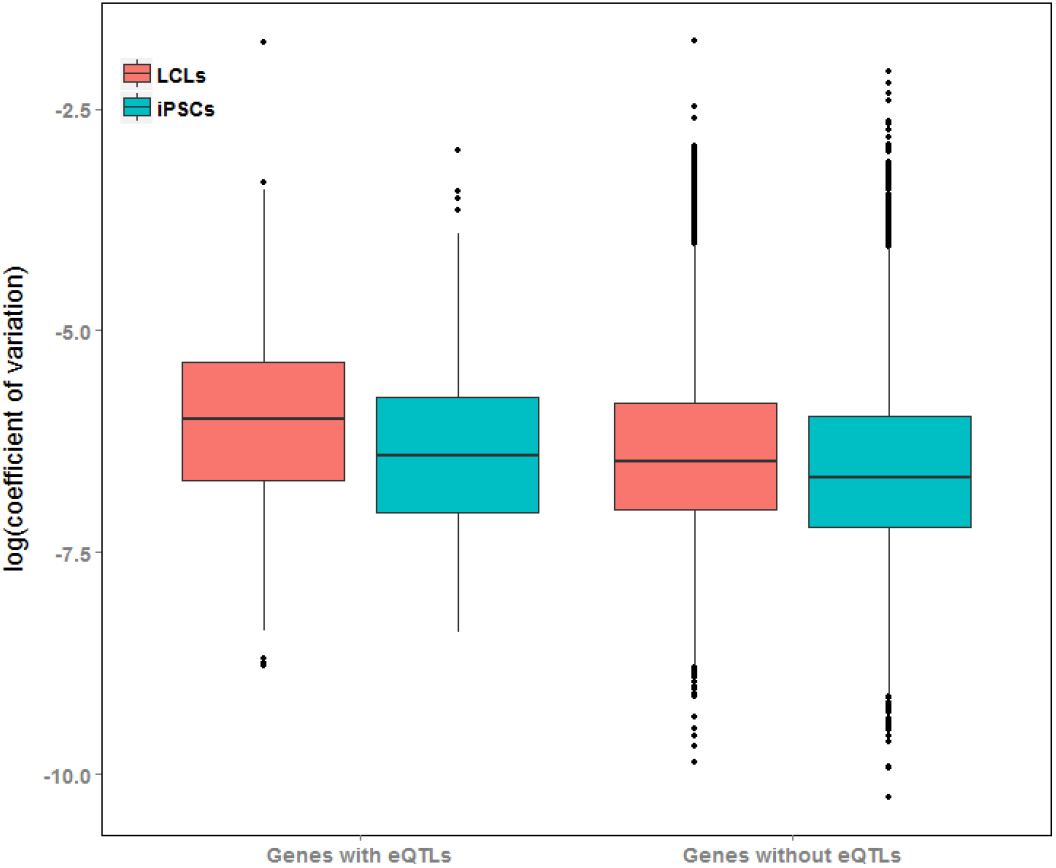
Genes with eQTLs are highly variable in both cell types. Boxplot of coefficients of variation of gene expression in genes with and without eQTLs previously identified in LCLs [6], plotted for LCLs and iPSCs.

## Discussion

The most useful renewable cell line models would retain a strong influence of individual of origin on their phenotypes, including molecular properties such as gene regulatory patterns. In previous work, we reported that freeze-thaw cycling of LCLs, a standard and required practice in long-term cell line maintenance, reduces the effect of donor on the cell line’s gene expression profile. In fact, we have found that whole-genome gene expression profiles from LCLs that were generated from different individuals are often as similar to each other as data from independently established replicates of LCLs from the same individual. We suggested that mature LCLs (those that have experienced one or more freeze-thaw cycle) may be clonally selected for, resulting in a convergent “LCL regulatory phenotype”, which has an advantage growing in culture but masks many of the original gene expression differences between the donor individuals. We noted that a subset of genes retained a high level of between-individual variation in their LCL gene expression profiles. Perhaps not surprisingly, these were enriched with genes for which eQTLs were identified in previous studies that considered gene expression data from mature LCLs.

Apart from the concern regarding the loss of much of the variation between donors, an intrinsic limitation of the LCL model system is that it theoretically represents the biology of only one primary cell type, B cells. In fact, existing collections of human population samples of renewable cell lines only include easily accessible primary tissues such as blood cells, adipocytes, and skin fibroblasts. Many cell types affected by disease, for example, cannot be directly studied using existing human cell line panels. In order to study variation in the most relevant phenotypes and disease processes, we need access to population samples that model additional cellular contexts.

The advent of iPSC technology may have provided the answer. It is now possible to establish renewable iPSC lines from population samples and differentiate them to multiple different cell types for which large collections are currently unavailable. One can establish iPSCs from fibroblast or fresh blood samples, but a most attractive possibility is to generate iPSC panels from the already available extensive collections of human LCLs. We thus asked whether individual variation in gene expression levels can be restored by reprograming LCLs into iPSC lines.

### Recovery of Individual Variation

We have shown that not only does the iPSC model exhibit a strong effect of donor on overall gene expression, but in fact the process of reprogramming highly manipulated immortalized cell lines to iPSCs recovers the inter-individual variation in gene expression lost during long term cell line maintenance.

The stronger clustering properties of expression data from iPSCs compared to LCLs suggest that iPSCs are better able to capture donor differences in gene regulation than LCLs. We could detect no significant difference between Euclidean distances within- and across-individuals in the projections of gene expression data from LCLs on the first two principal components of variation, indicating that donor is not a significant global source of gene expression variation in the LCL model. This observation is consistent with our previous findings [15]. In contrast, we observed a dramatic increase in the number of genes whose expression was significantly associated with donor in iPSCs, and a higher average variance in expression explained by individual of origin in the iPSCs compared with LCLs. These findings indicate that iPSCs reprogrammed from LCLs are a suitable model for studies of donor differences in gene regulation and genotype-phenotype interactions.

We note that despite their diminished ability to reflect donor differences, attempts to identify instances of genetic regulation of LCL gene expression in mature cell lines have been considered largely successful. Indeed, here we report that genes previously identified as associated with an eQTL in LCLs exhibit higher variance in mature LCLs than those without one. However, our observations suggest that, for future eQTL mapping studies, iPSCs may be a better system than LCLs. While 24% of genes with a significant donor effect in LCLs are associated with a previously identified eQTL in LCLs, only 5.6% of genes with a significant donor effect in iPSCs are associated with such an eQTL. Although expressed in LCLs, often at appreciable levels (S10 Fig.), the majority (>95%) of genes with a strong donor effect in iPSCs do not show such an effect in LCLs. Put together, these observations support our assertion that iPSC can be a better model than LCLs for detecting eQTLs, and more generally, for studies of inter-individual differences in gene regulation.

### Technical Noise Associated With Reprogramming

In any cell model, it is important to consider the magnitude of noise introduced by cell culture relative to biological signal. We note a substantial decrease in within-individual expression correlations for a cell line with a low PluriScore, indicating that we should perhaps reconsider acceptable scores for studies of individual phenotypic variation. However, other technical considerations do not seem to have a marked effect on overall clustering properties. For example, data from the single iPSC line that retained EBV (individual 3, replicate 2; S3 Fig.) clustered with the other iPSC lines derived from that individual. Additionally, we reprogrammed iPSCs in four groups and collected expression data at varying passages (between passage 11 and 13, S2 Table) without apparent batch effects. Because it is currently unclear which factors significantly affect our ability to detect donor differences, potential sources of noise need to be more systematically studied and appropriately controlled for.

Much of the excitement surrounding iPSCs is based on their ability to differentiate into terminal cell types, providing a renewable substitute for previously inaccessible tissues. Our study does not provide direct evidence that iPSC-derived differentiated cells will also reflect donor differences, however because the pluripotent state is relatively well-conserved compared to terminal cell types [27,28], we expect that tissues derived from iPSCs will demonstrate an even stronger donor effect on gene expression. That said, we suggest that this expectation needs to be independently confirmed in each differentiated cell type before they are carried into further studies.

### Conclusion

Because LCLs are available in large banks that represent panels of ethnic groups and disease populations, they are a popular cell model for genetic research and have been extensively studied. Recent advances in iPSC reprogramming protocols [21,22] have also positioned LCLs as a promising source of starting material for iPSC generation. Here, we have presented the recovery of donor gene expression patterns through the process of reprogramming highly manipulated LCLs to iPSCs, both validating the choice of iPSCs to study donor differences in physiology and the use of LCLs as an appropriate starting material for iPSC generation.

## Materials and Methods

### Sample Acquisition

Whole blood was collected from six healthy Caucasian donors by Research Blood Components LLC (Brighton, MA) with IRB consent between 2009 and 2010. B-Cell isolation and LCL generation were performed at the University of Chicago as described previously [14]. Between February 2011 and October 2012, each line was thawed, cultured, and re-frozen every three months, for a total of six freeze-thaw cycles prior to use in our study [15]. LCLs were cultured in RPMI with 20% FBS and frozen in Recovery Cell Culture Freezing Media (Life Technologies)

### iPSC Generation and Validation

All cell culture was performed at 37°C, 5% CO_2_, and atmospheric O_2_. From each individual, three biological replicates of LCLs were reprogrammed to iPSCs using a similar method to that described previously [21,22]. LCLs were transfected in four batches between August 2013 and January 2014 (S2 Table). One million cells were transfected with 2 μg of each episomal plasmid encoding *OCT3/4, shP53, Lin28, SOX2, L-MYC, KLF4,* and GFP(Addgene plasmids 27077, 27078, 27080, 27082 [20]) using the Amaxa transfection program X-005. For more details see: http://giladlab.uchicago.edu/data/LCL_Reprogramming.pdf. Transfected cells were grown in suspension for a week in hESC media (DMEM/F12 supplemented with 20% KOSR, 0.1mM NEAA, 2mM GlutaMAX, 1% Pen/Strep, 0.1 mM BME, and 12.5 ng/mL human bFGF) supplemented with 0.5mM sodium butyrate between days 2-12 post-nucleofection. After seven days, cells were plated on gelatin-coated plates with CF-1 irradiated mouse embryonic fibroblasts and manually passaged as colonies for at least 10 passages. After day 12, cells were grown in hESC media without sodium butyrate. Media was changed every 48 hours. Cell pellets were collected and stored at −80° C until extraction. One biological replicate from individual five failed to reach passage ten and was excluded from all analyses.

Embryoid body assays were performed following the protocol used by Romero et al [29]. Briefly, embryoid bodies were generated by manual colony detachment and were grown in suspension for seven days on low adherent plates in bFGF-free hESC media. They were then plated on 12 well gelatin-coated plates and grown for another seven days in DMEM-based media. Cells were fixed and stained using antibodies against nestin (1:250 SC-71665, Santa Cruz Biotech), α-smooth muscle actin (1:1500, CBL171, Millipore), alpha-Fetoprotein (1:100, SC-130302, Santa Cruz Biotech), and HNF3β (1:100 SC-6554, Santa Cruz Biotech) to detect ectoderm, mesoderm, and endoderm lineages respectively.

DNA was extracted using ZR-Duet DNA/RNA MiniPrep (Zymo) kits according to the manufacturer’s instructions. To assess for the presence of plasmid or EBV genome in iPSCs, PCR was performed using the genomic DNA collected from the iPSCs as template (collected at the same time as expression measurements) with primers designed to amplify the 3’ end of the EBNA-1 gene (present in both the EBV genome and all reprogramming plasmids) and NEBNext High-Fidelity 2X PCR Master Mix. For the sample with detectable EBNA-1, we also performed genomic PCR using primers to amplify a region common to all PXCLE reprogramming plasmids, and primers that amplify the BBRF1/LMP2 gene found only in the EBV genome to determine the source of foreign DNA. Primer sequences are available in S6 Table. Fibroblast DNA containing reprogramming plasmids at 0.02 pg/μL was used as a positive control for the PXCLE and EBNA-1 primer sets. LCL DNA (from YRI lines 18508 and 19238) were used as positive controls for the EBV and EBNA-1 primer sets. Fibroblast DNA was used as a negative control for all primer sets.

RNA was extracted using ZR-Duet DNA/RNA MiniPrep kits according to the manufacturer’s instructions with the addition of a DNAse treatment step prior to RNA extraction. cDNA was then synthesized using Maxima First Strand cDNA Synthesis Kit (Thermo-Scientific.) RT-PCR for endogenous transcripts of three pluripotency-related transcription factors was performed for all iPSC lines using SYBR Select master mix (Life Technologies.) Primers sequences are available in S6 Table. Data were analyzed using Viia7 software (Life Technologies). All expression levels were normalized to GAPDH. Expression was measured relative to a randomly selected iPSC line.

### Gene Expression Quantification

Cell pellets were obtained from LCLs immediately before transfection and from stable iPSCs after at least ten passages. RNA concentration and quality was estimated using the Agilent 2100 Bioanalyzer. Donor expression profiles were quantified using Illumina HumanHT-12 v4 Expression BeadChip Microarrays by the Functional Genomics Core at University of Chicago. Samples were hybridized across three array batches. Biological replicates from an individual were assigned to different batches to exclude a relationship between batch and individual. The array data were also used for the PluriTest assay as described previously [23].

### Data Processing and Analysis

Raw probe data were filtered for probes whose target transcripts were detected as expressed (*P* < 0.05) in at least two samples. Probes targeting expressed transcripts were then mapped to the hg19 reference genome and those that did not map uniquely to an Ensembl gene ID, or contained a HapMap SNP with MAF < 0.01 in CEU populations were excluded as described previously [15]. After filtering, probe intensities from all samples were background corrected, quantile-normalized, and log-2-transformed using the R package ‘*lumi*’ [30]. For genes represented by multiple probes, only the 3’ most probe was included in subsequent analyses to represent the most complete transcript. Finally, array batch was corrected for using an empirical Bayes method implemented in the R package *‘sva’[24,31]* This data is available in S1 Table.

Differential expression was estimated using a linear model based empirical bayes method implemented in the R package *‘limma [25]’*. Dendrograms were generated for matrices of pairwise Pearson product-moment correlation coefficients. For principal component analysis, expression data was mean-centered by gene across all individuals. The outlier individual 4-2 was omitted prior to hierarchical clustering analysis and PCA. All analyses, figures, and tables presented in the supplement include data from all individuals. Proportion of variance due to donor was estimated as the adjusted *R*^2^ value from a linear model including a term for each individual. Genes with FDR-adjusted p-values < 0.05 from a one-way ANOVA across individuals were classified as significantly associated with donor. eQTL data were downloaded from the Pritchard group eQTL browser: http://eqtl.uchicago.edu. Functional group enrichment was assessed using the web-based gene annotation database Lynx: http://lynx.ci.uchicago.edu using all expressed genes subjected to our filtering criteria as background.

## Acknowledgment

We thank Minal Çalişkan for providing the LCLs used in the study and all members of the Gilad lab for helpful discussion about data analysis and study design, with special thanks to Nicholas Banovich, John Blischak, Sidney Wang, and Irene Gallego Romero. This work was supported by grants HL092206 to YG and MH084703 to YG and JKP. JKP is also funded by Howard Hughes Medical Institute. Gene expression data are available at the GEO database, accession #GSE64263.

## Supporting Information Legends

**S. Figure 1: Embryoid body assay:** Immunocytochemistry approach to test for a cell line’s ability to spontaneously differentiate through endoderm: HNF3β and α-fetoprotein(AFP), mesoderm: smooth muscle actin (SMA), and ectoderm: nestin lineages. Scale bar: 200 μm. Individual channel levels, brightness, and contrast were adjusted using Adobe Photoshop CS6.

**S. Figure 2: PCR for EBNA-1:** Reprogramming vectors and Epstein-Barr virus are absent in all iPSC lines except 3-2 (see Figure S3)

**1**.4-1 iPSC, **2**. 6-3 iPSC, **3**. 5-2 iPSC, **4**. 3-3 iPSC, **5**. 3-2 iPSC, **6**. 6-2 iPSC, **7**. 2-3 iPSC, **8**. 5-1 iPSC, **9**. 1-2 iPSC, **10**. 1-1 iPSC, **11**. 3-1 iPSC, **12**. 3-3 iPSC, **13**. 2-2 iPSC, **14**. 2-1 iPSC, **15**. 4-2 iPSC, **16**. 1-3 iPSC, **17**. 6-1 iPSC, **18**. Reprogramming plasmids (positive control) **19**. LCL DNA (YRI lines 18508 and 19238, positive control). **20**. Fibroblast DNA (negative control), **21**. Water

**S. Figure 3: PCR for PCXLE (plasmid) and EBV genome for iPSC line 3-2:** iPSC 3-2 exhibits presence of EBV and absence of reprogramming plasmids.

**1**. 3-2 iPSC/EBNA-1 primer set, **2**. 3-2 iPSC/PXCLE primer set, **3**. 3-2 iPSC/EBV primer set, **4**. Reprogramming plasmid template/EBNA-1 primer set, **5**. Reprogramming plasmid template/PXCLE primer set, **6**. Reprogramming plasmid template/EBV primer set, **7**. LCL DNA/EBNA-1 primer set, **8**. LCL DNA/PCXLE primer set, **9**. LCL DNA/EBV primer set. **10**. Fibroblast DNA/EBNA-1 primer set **11**. Fibroblast DNA/PCXLE primer set **12**. Fibroblast DNA/EBV primer set.

**S. Figure 4: Clustering Analysis:** A. Results from hierarchical clustering analysis of microarray gene expression and expression data projections on principal components axes 1 and 2 from cycle 7 LCLs and B. iPSCs. Includes data from all lines.

**S. Figure 5: Correlation Heatmaps:** Heatmap generated from pairwise correlation matrix (pearson product-moment correlation coefficients) for A. LCLs and B. iPSCs. Includes data from all lines.

**S. Figure 6: Within and between individual expression correlation:** Pairwise Pearson correlation coefficients for gene expression data from lines derived from the same individual and across different individuals for both cell types. iPSCs demonstrate increased correlation both within and across individuals compared with LCLs. Includes data from all lines.

**S. Figure 7: Density plots of gene expression variance:** A. Density plot of between donor variance to within donor variance in gene expression for all expressed genes in iPSCs and LCLs including data from all lines. The dotted line indicates the threshold corresponding with significant association between gene expression and individual of origin. X-axis was truncated at 8.0; 0.67% of the data are not shown. B. Density plot of total variance in LCLs and iPSCs including data from all lines. X-axis was truncated at 0.1; 3.7% of the data are not shown.

**S. Figure 8: Unadjusted p-values for donor effect:** Histogram of unadjusted p-values from ANOVA F-test across the factor individual of origin for A. iPSCs and B. LCLs. Includes data from all lines.

**S. Figure 9: Coefficient of variation in genes with and without eQTLs:** Boxplot of coefficients of variation of gene expression in genes with and without eQTLs previously identified in LCLs plotted for LCLs (*P* < 10^-10^) and iPSCs (*P* = 0.01). Includes data from all lines.

**S. Figure 10: Expression in genes with a donor effect in iPSCs:** Mean gene expression levels for genes for which a donor effect was detected in iPSCs compared to all genes. Genes with a strong iPSC donor effect are expressed in LCLs, in fact with a higher mean expression value than the genome-wide average (*P* < 10^-15^). These genes exhibit significantly higher expression than the genome-wide average in iPSCs as well (*P* < 10^-15^)

**S. Table 1: Expression data and analysis results for all 15,306 expressed genes**. Columns 1:34: processed expression levels (see methods) for each sample, 35: p-value from test for differential expression between LCLs and iPSCs (limma) 36: FDR-adjusted p-value for differential expression between LCLs and iPSCs (limma). 37: FDR-adjusted F-test p-values from test for donor effect in iPSCs. 38: FDR-adjusted F-test p-values from test for donor effect in LCLs.

**S. Table 2: Sample information:** Includes the following information for all lines: sample ID, gender, reprogramming, extraction, and array batch assignments and dates, RIN scores, passage of RNA collection for iPSCs, and PluriTest scores for all samples.

**S. Table 3: Euclidean distances between clusters in principal components analysis:** Euclidean distances of sample projections on the first two principle component axes within and between individuals averaged over all samples for each cell type, demonstrating significantly lower within-individual distances compared to between-individual distances in iPSCs but not LCLs, regardless of outlier inclusion status.

**S. Table 4: Lynx enrichment analysis in LCLs:** Tissue and disease enrichment results for genes with a significant donor effect in LCLs at an FDR-cutoff of 0.05. Data downloaded from the online database Lynx.

**S. Table 5: Lynx enrichment analysis in iPSCs:** Tissue and disease enrichment results for genes with a significant donor effect in iPSCs at an FDR-cutoff of 0.05. Data downloaded from the online database Lynx.

**S. Table 6: PCR primer information:** Sequence, use, and source of primers used for PCR and qPCR.

## References

1. Hu VW, Frank BC, Heine S, Lee NH, Quackenbush J (2006) Gene expression profiling of lymphoblastoid cell lines from monozygotic twins discordant in severity of autism reveals differential regulation of neurologically relevant genes. BMC Genomics 7: 118.

2. Huang RS, Duan S, Kistner EO, Hartford CM, Dolan ME (2008) Genetic variants associated with carboplatin-induced cytotoxicity in cell lines derived from Africans. Molecular cancer therapeutics 7: 3038–3046.

3. Wen Y, Gamazon ER, Bleibel WK, Wing C, Mi S, et al. (2012) An eQTL-based method identifies CTTN and ZMAT3 as pemetrexed susceptibility markers. Hum Mol Genet 21: 1470–1480.

4. Ziliak D, O'Donnell PH, Im HK, Gamazon ER, Chen P, et al. (2011) Germline polymorphisms discovered via a cell-based, genome-wide approach predict platinum response in head and neck cancers. Transl Res 157: 265–272.

5. Moyer AM, Fridley BL, Jenkins GD, Batzler AJ, Pelleymounter LL, et al. (2011) Acetaminophen-NAPQI hepatotoxicity: a cell line model system genome-wide association study. Toxicological sciences : an official journal of the Society of Toxicology 120: 33–41.

6. Pickrell JK, Marioni JC, Pai AA, Degner JF, Engelhardt BE, et al. (2010) Understanding mechanisms underlying human gene expression variation with RNA sequencing. Nature 464: 768–772.

7. Banovich NE, Lan X, McVicker G, van de Geijn B, Degner JF, et al. (2014) Methylation QTLs Are Associated with Coordinated Changes in Transcription Factor Binding, Histone Modifications, and Gene Expression Levels. PLoS Genet 10: e1004663.

8. Choy E, Yelensky R, Bonakdar S, Plenge RM, Saxena R, et al. (2008) Genetic analysis of human traits in vitro: drug response and gene expression in lymphoblastoid cell lines. PLoS Genet 4: e1000287.

9. Plagnol V, Uz E, Wallace C, Stevens H, Clayton D, et al. (2008) Extreme clonality in lymphoblastoid cell lines with implications for allele specific expression analyses. PLoS One 3: e2966.

10. Stark AL, Zhang W, Mi S, Duan S, O'Donnell PH, et al. (2010) Heritable and non-genetic factors as variables of pharmacologic phenotypes in lymphoblastoid cell lines. Pharmacogenomics J 10: 505–512.

11. Hannula K, Lipsanen-Nyman M, Scherer SW, Holmberg C, Höglund P, et al. (2001) Maternal and paternal chromosomes 7 show differential methylation of many genes in lymphoblast DNA. Genomics 73: 1–9.

12. Carter KL, Cahir-McFarland E, Kieff E (2002) Epstein-barr virus-induced changes in B-lymphocyte gene expression. J Virol 76: 10427–10436.

13. Min JL, Barrett A, Watts T, Pettersson FH, Lockstone HE, et al. (2010) Variability of gene expression profiles in human blood and lymphoblastoid cell lines. BMC Genomics 11: 96.

14. Caliskan M, Cusanovich DA, Ober C, Gilad Y (2011) The effects of EBV transformation on gene expression levels and methylation profiles. Hum Mol Genet 20: 1643–1652.

15. Calışkan M, Pritchard JK, Ober C, Gilad Y (2014) The effect of freeze-thaw cycles on gene expression levels in lymphoblastoid cell lines. PLoS One 9: e107166.

16. Mills JA, Wang K, Paluru P, Ying L, Lu L, et al. (2013) Clonal genetic and hematopoietic heterogeneity among human-induced pluripotent stem cell lines. Blood 122: 2047–2051.

17. Boulting GL, Kiskinis E, Croft GF, Amoroso MW, Oakley DH, et al. (2011) A functionally characterized test set of human induced pluripotent stem cells. Nat Biotechnol 29: 279–286.

18. Kajiwara M, Aoi T, Okita K, Takahashi R, Inoue H, et al. (2012) Correction for Kajiwara et al., Donor-dependent variations in hepatic differentiation from human-induced pluripotent stem cells. Proceedings of the National Academy of Sciences 109: 14716–14716.

19. Rouhani F, Kumasaka N, de Brito MC, Bradley A, Vallier L, et al. (2014) Genetic background drives transcriptional variation in human induced pluripotent stem cells. PLoS Genet 10: e1004432.

20. Okita K, Matsumura Y, Sato Y, Okada A, Morizane A, et al. (2011) A more efficient method to generate integration-free human iPS cells. Nat Methods 8: 409–412.

21. Choi SM, Liu H, Chaudhari P, Kim Y, Cheng L, et al. (2011) Reprogramming of EBV-immortalized B-lymphocyte cell lines into induced pluripotent stem cells. Blood 118: 1801–1805.

22. Rajesh D, Dickerson SJ, Yu J, Brown ME, Thomson JA, et al. (2011) Human lymphoblastoid B-cell lines reprogrammed to EBV-free induced pluripotent stem cells. Blood 118: 1797–1800.

23. Müller FJ, Schuldt BM, Williams R, Mason D, Altun G, et al. (2011) A bioinformatic assay for pluripotency in human cells. Nat Methods 8: 315–317.

24. Johnson WE, Li C, Rabinovic A (2007) Adjusting batch effects in microarray expression data using empirical Bayes methods. Biostatistics 8: 118–127.

25. Smyth GK (2004) Linear models and empirical bayes methods for assessing differential expression in microarray experiments. Stat Appl Genet Mol Biol 3: Article3.

26. Sulakhe D, Balasubramanian S, Xie B, Feng B, Taylor A, et al. (2014) Lynx: a database and knowledge extraction engine for integrative medicine. Nucleic Acids Res 42: D1007–1012.

27. Garfield DA, Runcie DE, Babbitt CC, Haygood R, Nielsen WJ, et al. (2013) The impact of gene expression variation on the robustness and evolvability of a developmental gene regulatory network. PLoS Biol 11: e1001696.

28. Roux J, Robinson-Rechavi M (2008) Developmental constraints on vertebrate genome evolution. PLoS Genet 4: e1000311.

29. Gallego Romero I, Pavlovic BJ, Hernando-Herraez I, Banovich NE, Kagan CL, et al. (2014) Generation of a Panel of Induced Pluripotent Stem Cells From Chimpanzees: a Resource for Comparative Functional Genomics. bioRxiv doi: 10.1101/008862.

30. Du P, Kibbe WA, Lin SM (2008) lumi: a pipeline for processing Illumina microarray. Bioinformatics 24: 1547–1548.

31. Leek JT, Johnson WE, Parker HS, Jaffe AE, Storey JD (2012) The sva package for removing batch effects and other unwanted variation in high-throughput experiments. Bioinformatics 28: 882–883.

